# Binge alcohol drinking alters the differential control of cholinergic interneurons over nucleus accumbens D1 and D2 medium spiny neurons

**DOI:** 10.1101/2022.06.22.497229

**Authors:** Jenya Kolpakova, Vincent van der Vinne, Pablo Gimenez-Gomez, Timmy Le, Gilles E. Martin

**Affiliations:** Brudnick Neuropsychiatric Research Institute, Department of Neurobiology, University of Massachusetts Chan Medical School, Worcester MA 01655, USA; Biology Department, Williams College, Williamstown MA 01247, USA; Graduate Program in Neuroscience, Morningside Graduate School of Biomedical Sciences, UMass Chan Medical School

**Author notes:** **Corresponding author** Gilles E. Martin. **Author contributions** JK, PGG, TL and GEM did the experiments, JK and VvdV analyzed data, JK, VvdV and GEM wrote the manuscript.

## Abstract

**Background:** Ventral striatal cholinergic interneurons (ChIs) play a central role in basal ganglia function by regulating associative learning and reward processing. In the nucleus accumbens (NAc), ChIs regulate glutamatergic, dopaminergic, and GABAergic neurotransmission. However, it is unclear how ChIs orchestrate the control of these neurotransmitters to determine the excitability of medium spiny neurons (MSNs) expressing either dopamine D1 or D2 receptors. Additionally, the effects of binge alcohol drinking on ChIs-mediated modulation of glutamatergic synaptic transmission in NAc MSNs are also undefined.

**Methods:** We optogenetically stimulated ChIs while recording evoked and spontaneous excitatory postsynaptic currents (sEPSCs) in D1- and D2-MSN of ChAT.ChR2.eYFPxDrd1.tdtomato mice. To determine the effect of ChIs on mouse behavior and alcohol consumption, we implanted ChAT.ChR2.eYFP mice with fiber optic cannulas and stimulated ChIs while mice were allowed to drink 20% alcohol using the binge alcohol drinking-in-the-dark (DID) paradigm.

**Results:** We demonstrated that NAc ChIs decrease the frequency of spontaneous excitatory postsynaptic currents (sEPSCs) in both D1- and D2-MSNs,. While inhibition of D1-MSNs glutamate release by ChIs depends on dopamine release, that of D2-MSNs results from a direct interactions between ChIs and glutamatergic terminals. Interestingly, after two weeks of binge alcohol drinking, the effect of ChIs stimulation on glutamate release was reversed in D1-MSNs, while its effect on D2-MSNs remained unchanged. Finally, *in vivo* optogenetic stimulation of NAc ChIs significantly increased alcohol consumption.

**Conclusions:** These results identify ChIs as a key target for the regulation of NAc circuitry and as a potential treatment of alcohol addiction.

## Introduction

Addiction is a disorder of the reward system (1, 2) where drugs of abuse distort the response to natural reinforcers leading to continued drug use, which, in turn, impairs brain function by interfering with the capacity to exert self-control over drug-taking behaviors such as binge drinking (1, 2). Binge alcohol drinking is the main mode of alcohol consumption in late adolescents and young adults and often serves as a gateway to alcohol dependence later in life (3). One of the main brain areas controlling drug taking behaviors is the nucleus accumbens (NAc), a forebrain region that encodes association between temporally unpredictable stimuli and the appropriate action to maximize reward or avoid punishment (4). MSNs expressing dopamine-1 and 2 receptors (D1- and D2-MSNs) are the sole output neurons of the NAc. D1- and D2-MSNs promote and inhibit reinforcement behavior, respectively(5), and optogenetic manipulation of their excitability has been causally linked to reward-seeking behaviors (6, 7). The role traditionally attributed to MSNs is that of integrators that receive a range of different inputs (glutamate, dopamine, acetylcholine, and GABA) from across the brain and determine the optimal behavioral response (7–9). In recent years, this view has been challenged by the observations that the integration of different inputs is mainly performed by a different cell population in the NAc: Cholinergic interneurons (ChIs) (10, 11).

Cholinergic interneurons (ChIs) make up only 1-2% of all neurons in the striatum(12), but play an outsize role in regulating NAc GABAergic (13), glutamatergic (14, 15) and dopaminergic synaptic transmission (16, 17) through their extensive projections (11). NAc ChIs generate unique bidirectional outcome responses during reward-based learning, signaling both positive (reward) and negative (reward omission) outcomes (18). Cholinergic receptor signaling has been shown to alter alcohol and other drugs’ consumption (19–21). Currently, both the role played by ChIs in orchestrating dopamine (DA) and glutamatergic synaptic transmission to regulate D1- and D2- MSNs excitability, as well as how alcohol exposure modulates this connection remain to be elucidated. Here we combine *in vitro* patch clamp, fast scan cyclic voltammetry (FSCV), optogenetics and behavioral recordings to answer these questions.

In alcohol-naïve mice, we demonstrate that optogenetic stimulation of ChIs decreases the frequency of spontaneous excitatory post-synaptic currents (sEPSCs), presumably through a presynaptic mechanism, in both D1- and D2-MSNs. In D1-MSNs, inhibition of glutamatergic synaptic transmission by ChIs is mediated by dopaminergic and cholinergic (nAChR and mAChR) receptors. Although ChIs induced a similar effect on sEPSPCs in D2-MSNs, this effect did not require DA. Instead, glutamatergic inhibition likely resulted from ChIs synapsing directly on glutamatergic terminals. Importantly, binge alcohol drinking differentially altered ChIs control of glutamatergic synaptic transmission in D1- and D2-MSNs. While the ChI-mediated decrease of sEPSCs frequency in D2-MSNs was unaffected, the ChI-induced inhibition of glutamatergic transmission in D1-MSNs seen in naïve mice was reversed and optogenetic stimulation became potentiating following preceding alcohol exposure. Finally, optogenetic stimulation of ChIs *in vivo* significantly increased alcohol consumption in mice, while not altering locomotion or saccharine and water consumption. Our findings elucidate mechanisms by which ChIs differentially control excitability of D1- and D2-MSNs in naïve and alcohol conditions, and their influence on binge alcohol drinking.

## Materials and Methods

A detailed description of all materials and methods can be found in Supplemental Methods and Materials. We used a binge alcohol drinking protocol in 6 – 10 week old mice as previously described (22, 23). Twenty-four hours after the last alcohol-drinking session, animals were sacrificed and 200 µm coronal sections containing the core nucleus accumbens were prepared. The external solution contained (in mmol/L): 126 NaCl, 2.5 KCl, 1.25 NaH_2_PO_4_.H_2_O, 1 MgCl_2_.H_2_O, 2 CaCl_2_.H_2_O, 26 NaHCO_3_, 10 D-Glucose, at room temperature. The pipette solution contained in mM: 120 K-methanesulfonate; 20 KCl; 10 HEPES; 2 ATP, 1 GTP, and 12 phosphocreatine. All recordings were performed at MSNs’ resting membrane potential around - 84 mV. 4-week old mice were anesthetized and implanted with an optic fiber cannula (Doric Lenses, CA, United States) located above the NAc (AP +1.5, ML ±1.5, DV −4.0 mm from Bregma).

## Results

ChAT.ChR2.eYFP and DrD1.TdTomato mouse lines were crossed to generate brain slices in which D1-MSNs could be identified while the role of ChIs in regulating glutamatergic synaptic transmission onto NAc MSNs could be assessed through optogenetic stimulation. Immunostaining showed the presence of eYFP and TdTomato reporters for ChIs and D1-MSNs, respectively, in the NAc (Fig. 1A). eYFP-positive neurons were confirmed to be ChIs by injecting incremental current steps and recording voltage responses: current-voltage relationships presented the hallmarks of cholinergic interneurons, i.e., depolarized resting membrane potential (∼-50 mV), large membrane resistance and sag, and spontaneous firing (Fig. 1B). To verify that ChIs expressed functional Channelrhodopsin (Fig. 1C), ChIs were stimulated with blue light (five light pulses at 20 Hz every 20 seconds for 2 minutes). This pattern faithfully evoked action potentials in all (n = 8) neurons tested (Fig. 1D).

**Figure 1:**
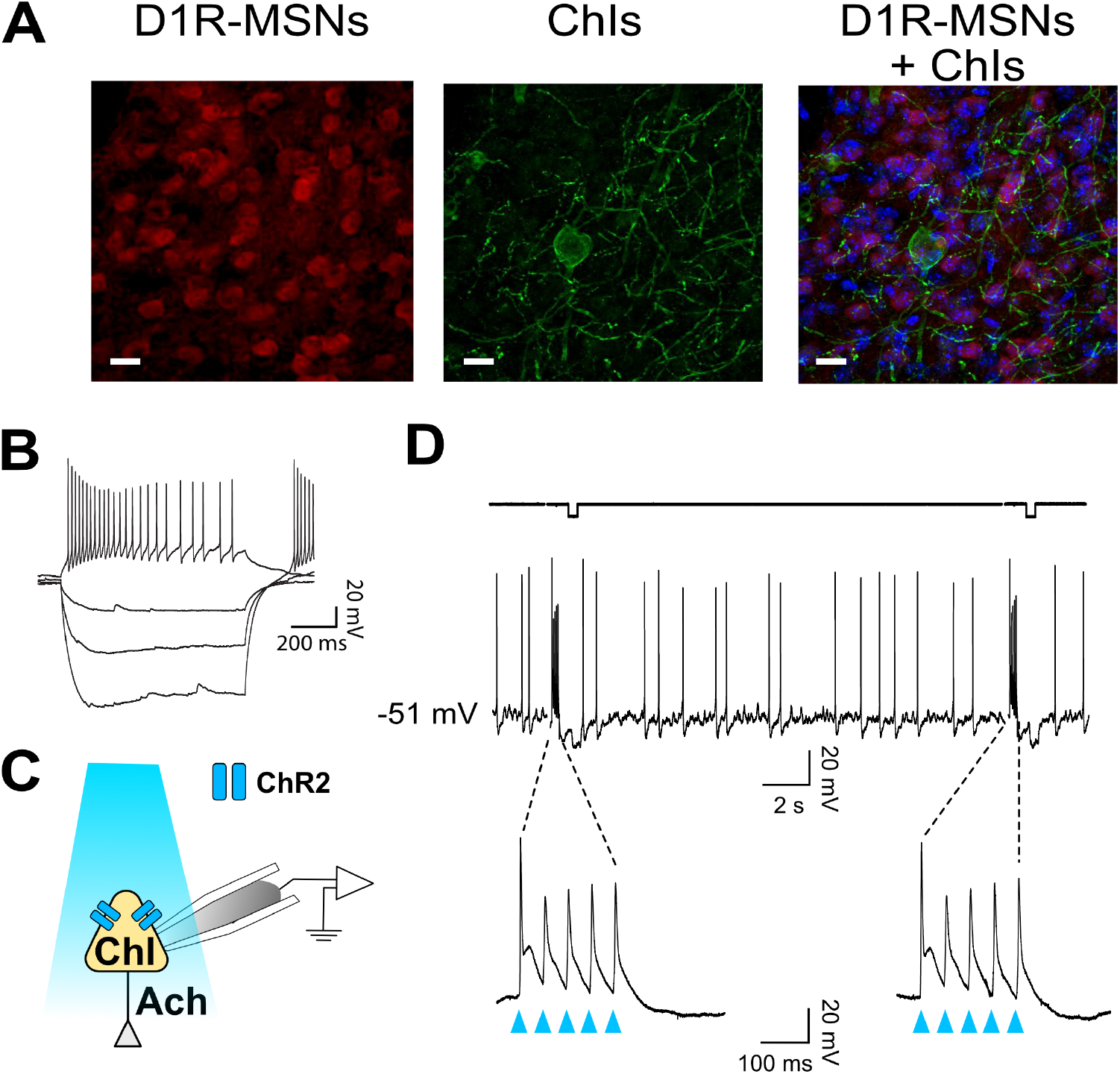
ChAT-ChR2-eYFP x DrD1-tdTomato mouse line to optogenetically stimulate ChIs and differentiate core NAc D1- and D2-MSNs. **A**. Immunostaining in ChAT-ChR2-eYFP x DrD1-tdTomato mouse line of D1-MSNs (left panel, red-fluorescence) and cholinergic interneurons (middle panel, eYFP fluorescence) in the nucleus accumbens. Right panel shows overlaid left and middle panels. All neurons are stained in blue. Scale bar 20 um. **B**. Representative voltage traces in response to incremental current steps (-150 pA to 0 in steps of 50) in a cholinergic interneuron. **C**. Schematic of ChI recording during optogenetic stimulation with blue light. **D**. Optogenetic stimulation (i.e., burst of five 20 Hz pulses every 20 seconds for 2 minutes) evokes action potentials in ChI.

### Cholinergic interneurons decrease glutamate release in D1- and D2-MSNs

To determine whether ChIs controlled glutamate release in D1-MSNs, sEPSCs were recorded in TdTomato-labeled neurons while ChIs were simultaneously stimulated (Fig. 2A) with a pattern described in Fig. 1. Current-voltage relationships confirmed that all recorded red epifluorescent neurons (Fig. 2Biii) were MSNs (Fig. 2Bi,ii). Often, the cell body of ChIs could be detected in the vicinity of recorded MSNs (Fig. 2Biv). sEPSCs were recorded at MSNs’ resting membrane potential (-85 ± 0.7 mV in a random sample of 10 neurons) for 4 minutes (Pre-stim; Fig. 2Ci) before stimulating ChIs for 2 minutes (Fig. 2Ci, blue arrowheads), followed by recording sEPSCs for 4 minutes (Post-stim; Fig. 2Ci). The inter-event intervals (IEIs) between sEPSCs lengthened during the Post-stim versus the Pre-stim interval (i.e., a decreased frequency; Fig. 2C, t(13) = 2.868, p = 0.0132, paired t-test, n = 14). Interestingly, ChI stimulation did not affect the amplitude of D1-MSN sEPSCs in ChAT-ChR2 mice (Fig. S1A, t(13) = 2.04, p = 0.0619, paired t-test, n = 14). To verify that these effects are specifically due to optogenetic ChI stimulation, D1- MSN sEPSCs were recorded in slices obtained from the DrD1.Tdtomato mouse line (Fig. 2Di). These recordings demonstrating that optical stimulation did not significantly affect IEIs (Fig. 2Dii, t(9) = 0.2512, p = 0.8073, paired t-test, n = 10) or amplitude (Fig. S1B, t(9) = 1.571, p = 0.1506, paired t-test, n = 10). Comparisons of IEI cumulative frequency distributions revealed a significant difference between Pre- and Post-ChI stimulation conditions in D1-MSNs ChAT.ChR2 mice (Fig. 2E, D = 0.07492, p <0.0001, K-S test) but not in D1-MSN TdTomato control mice (Fig. 2F, D = 0.03261, p = 0.0603, K-S test). Next, the relationship between sEPSCs’ IEI size and the ChI-mediated effect was assessed. To address this relationship, sets of 25 consecutive IEIs were ordered by IEI and the 50 (median), 25 and 75 (shoulders), as well as the 5 and 95 (extremes) percentile values were analyzed^72^. No effect of different IEI size distribution on ChI-mediated IEI increase was observed (Fig. 2E inset, F(4, 907) = 0.3362, p = 0.85, Mixed-model general linear modeling (MM GLM)), indicating that EPSCs are uniformly altered by ChIs. Finally, a significant difference in response to ChI stimulation was observed between ChR2 and TdTomato groups (Fig. 2G; F(1,2265) = 25.58, p < 0.0001, MM GLM, with Tukey Post-hoc tests revealing significant increases in IEI size between Pre- and Post-stimulation in ChR2 but not in TdTomato controls). Interestingly, ChI stimulation had no effects on electrically evoked EPSPs in D1-MSNs (Fig. S1C, F(2,10) = 0.973, p = 0.3785, RM one-way ANOVA) and TdTomato control mice (Fig. S1D, F(2,6) = 1.7, p = 0.238, RM one-way ANOVA). These results indicate that optogenetic stimulation of ChIs decreases only spontaneous glutamate release onto D1-MSNs, a likely presynaptic effect.

**Figure 2:**
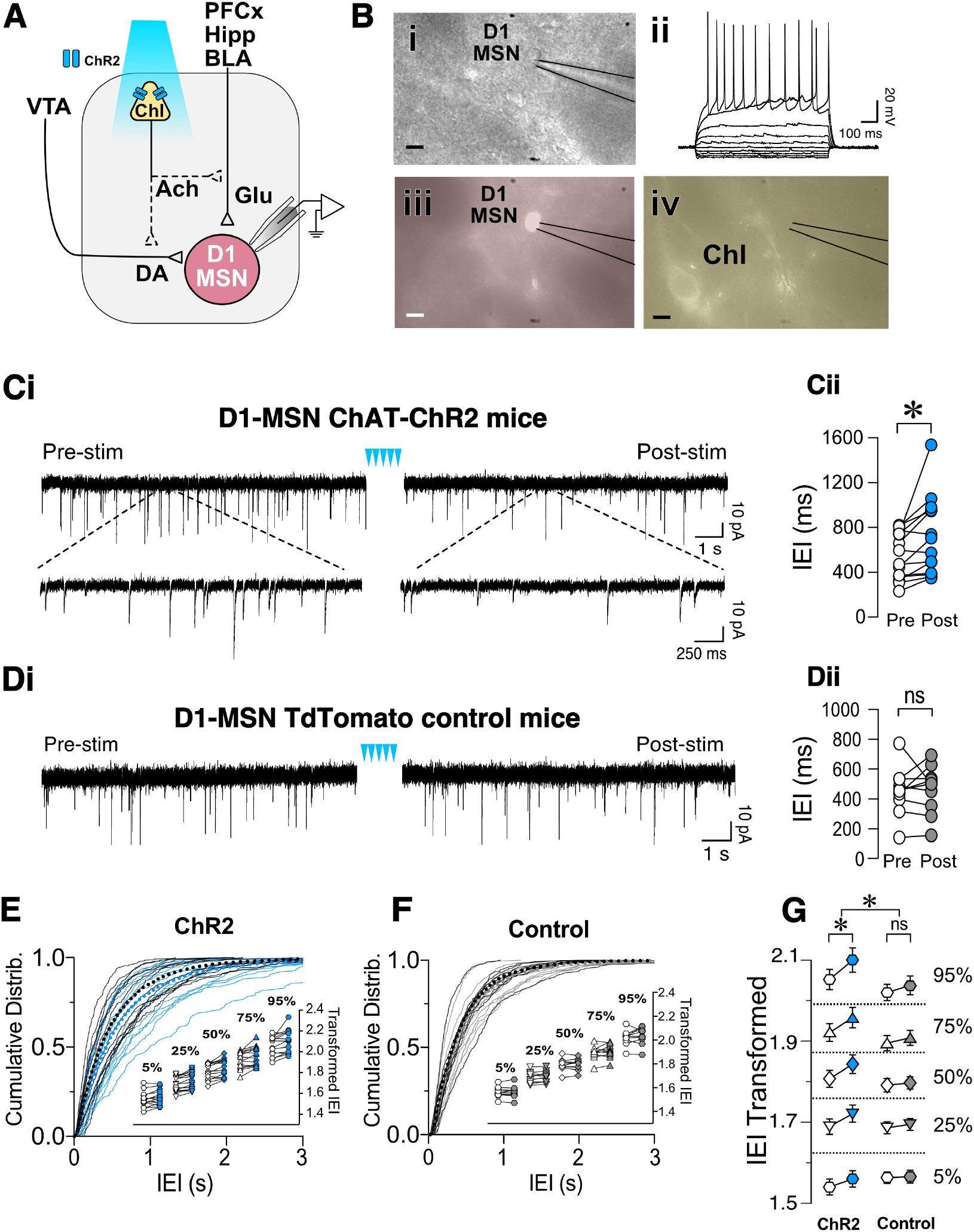
Optogenetic stimulation of ChIs decreases sEPSCs frequency in D1-MSNs. **A**. Schematic of the experimental setup: whole cell recording of D1-MSNs sEPSC during optogenetic stimulation of ChIs. **B**. Same slice images of representative MSNs (i, DIC), corresponding IV traces to confirm MSN identity (ii), red-epifluorescence to verify the cell is D1R+ (iii), and eYFP epifluorescence of ChI cell in proximity of the recording (iv). Scale bar 10 um. **Ci**. Representative sEPSCs in D1-MSNs in ChAT-ChR2 mice before (Pre) and after (Post) ChI optogenetic stimulation (blue arrowheads). **Cii**. Average sEPSCs inter-event intervals (IEI) in Pre (white circles) and Post (blue circles) ChI optogenetic stimulation in ChAT-ChR2 D1-MSNs (n = 14). **Di**. Representative traces of sEPSCs in D1-MSNs of tdTomato control mice before (Pre) and after (Post) ChI stimulation. **Dii**. Average EPSCs inter-event intervals (IEI) in Pre (white circles) and Post (gray circles) ChI optogenetic stimulation in control tdTomato D1-MSNs (n = 10). **E:** Cumulative frequency distribution of D1-MSN sEPSCs IEI in ChAT-ChR2 mice group Pre (black traces) and Post (blue traces) ChI optogenetic stimulation. Each solid line represents a neuron. Average traces for Pre and Post conditions are shown in dotted black and blues lines, respectively. **Inset**. Cumulative distributions of ChAT-ChR2 D1-MSNs EPSCs IEIs broken into percentiles of distribution to quantify median (50%), shoulders (25 and 75%), and extreme values (5 and 94%) of distribution, that are 1^10 transformed to normalize the distribution. **F:** Cumulative frequency distribution of D1-MSNs inter-event intervals (IEI) of sEPSCs before (Pre, black lines) and after (Post, gray lines) ChI optogenetic stimulation in tdTomato control mice. Each solid line represents a neuron. Average traces are shown in dotted black and blues lines for Pre and Post conditions, respectively. **Inset**. Same as inset in E, but in tdTomato D1-MSN controls. **G:** Percentiles of cumulative distribution of transformed IEIs of EPSCs in ChAT-ChR2 D1-MSNs (Pre, white circles, Post, blue circles) and control TdTomato D1-MSNs (Pre, white circles, Post, gray circles). *p < 0.05, ns: no significant difference

To determine whether ChIs similarly regulated glutamatergic synaptic transmission in putative D2-MSNs, non-fluorescent MSNs were recorded while stimulating ChIs optogenetically (Fig. 3A and B). Current-voltage relationships confirmed that all recorded neurons were MSNs (Fig. 3Bii). As with D1-MSNs, putative D2-MSNs’ baseline sEPSCs was recorded for 4 minutes before (Pre) and after (Post) optogenetic ChIs stimulation. Similar to D1-MSNs, blue light stimulation significantly increased average IEIs (i.e., decrease frequency; Fig. 3C, t(11) = 2.27, p = 0.0443, paired t-test, n = 12) in sliced obtained from ChAT-ChR2 mice but not in TdTomato control slices (Fig. 3D, t(8) = 0.604, p = 0.563, paired t-test, n = 9). Likewise, significant increase of IEI sEPSCs cumulative frequency distribution in Pre vs Post groups in D2-MSNs of ChAT- ChR2 mice was observed (Fig. 3E, D = 0.048, p = 0.0011, K-S test, n = 12), but not in D2-MSNs of TdTomato control mice (Fig. 3F, D = 0.0175, p = 0.7278, K-S test, n = 9). As with D1-MSNs, significant increases of IEI were observed at all percentiles in D2-MSNs of ChAT-ChR2 (Fig. 3E inset), but not TdTomato control mice (Fig. 3F, inset). Finally, a significant difference in response to ChI stimulation between ChR2 and TdTomato control groups was observed (Fig. 3G; F(1,1915) = 18.05, p<0.0001, MM GLM, with Tukey post-hoc tests showing significant increase in IEI duration between Pre- and Post-intervals in ChR2 but not in TdTomato controls). No effects of ChI stimulation on D2R MSNs sEPSC amplitude was observed in ChR2 (Fig. S2A, t(11) = 0.129, p = 0.8995, paired t-test), or TdTomato control group (Fig. S2B, t(8) = 0.277, p = 0.789, paired t-test). Interestingly, electrically evoked EPSPs in D2-MSNs following ChI optogenetic stimulation had a significantly decreased amplitude in ChR2 groups (Fig. S2C, F(2,8) = 6.01, p = 0.0145), but not in TdTomato control groups (Fig. S2D, F(2,6) = 0.20, p = 0.742). These results indicate that ChIs stimulation decreases glutamate release presynaptically onto D1- and D2-MSNs.

**Figure 3:**
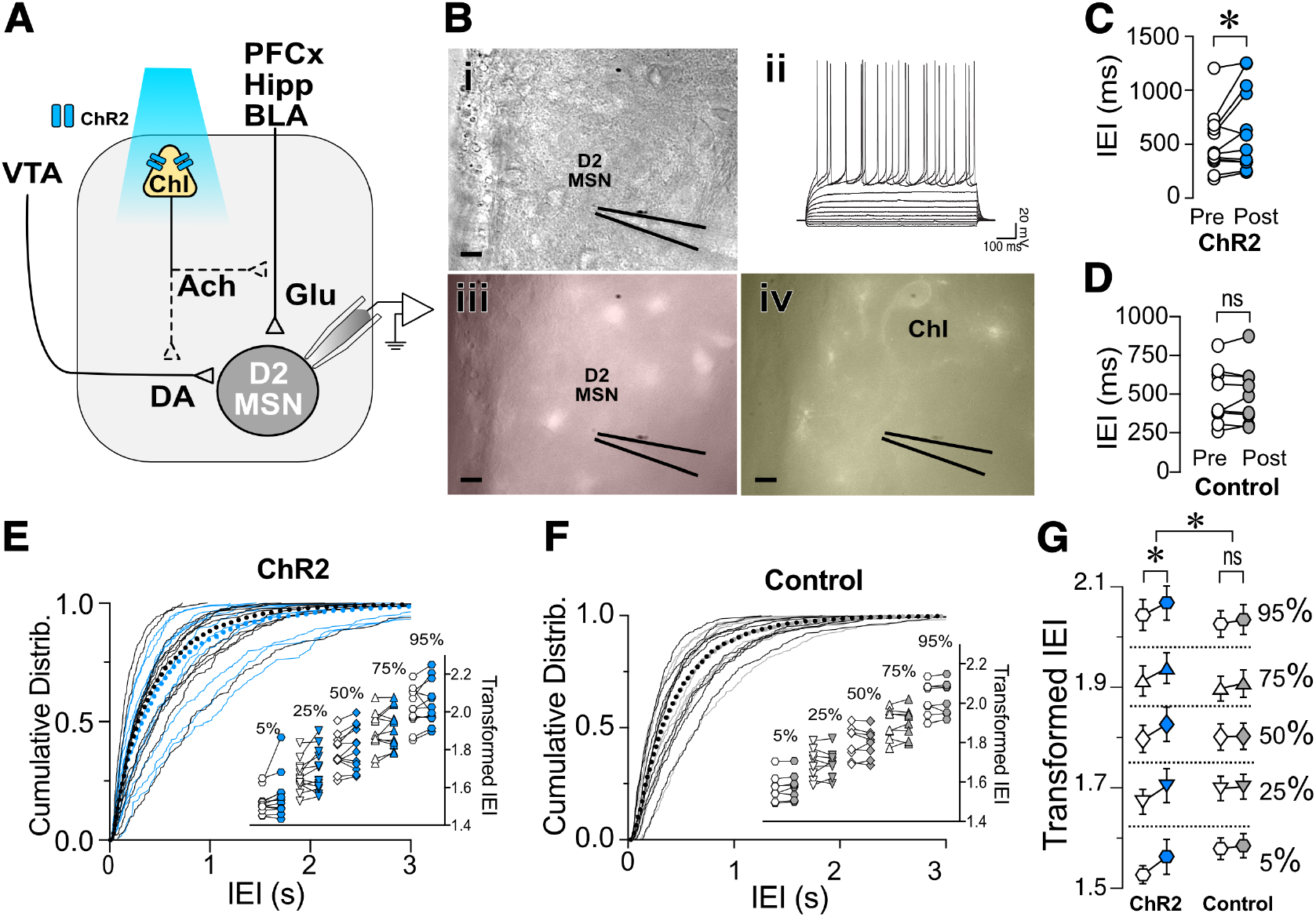
Optogenetic stimulation of ChIs decreases sEPSCs frequency in D2-MSNs. **A**. Schematic of the experimental setup: whole cell sEPSC recording of D2-MSNs during optogenetic stimulation of ChIs. **B**. Same slice images of representative MSNs (i, DIC), corresponding IV traces to confirm MSN identity (ii), lack of red-epifluorescence to verify the cell is D1R(-) (iii), and eYFP epifluorescence of ChI cell in the proximity of the recording (iv). Scale bar 10 um. **C**. Average sEPSCs IEI before (Pre, white circles) and after (Post, blue circles) optogenetic stimulation of ChIs in ChAT-ChR2 D2-MSNs (n = 12). **D**. Average sEPSCs IEI before (Pre, white circles) and after (Post, gray circles) optogenetic stimulation of ChIs in tdTomato control mice (n = 9). **E**. Cumulative frequency distribution of D2-MSN IEI of sEPSCs in ChAT-ChR2 mice before (Pre, black traces) and after (Post, blue traces) ChI optogenetic stimulation. Each solid line represents a neuron. Average traces are shown in dotted black and blues lines for Pre and Post conditions, respectively. **Inset**. Cumulative distributions of ChAT-ChR2 D2-MSNs EPSCs IEIs broken into percentiles of distribution to quantify median (50%), shoulders (25 and 75%), and extreme values (5 and 94%) of distribution, that are 1^10 transformed to normalize the distribution. **F**. Cumulative frequency distribution of D2-MSN IEI of sEPSCs in tdTomato mice before (Pre, black traces) and after (Post, blue traces) ChI optogenetic stimulation. Each solid line represents a neuron. Average traces are shown in dotted black and blues lines for Pre and Post conditions, respectively. **Inset**. Same as inset in E, but in TdTomato D2-MSN controls. **G**. Percentiles of cumulative distribution of transformed IEIs of EPSCs in ChAT-ChR2 D2-MSNs (Pre, white circles, Post, blue circles) and control TdTomato D2-MSNs (Pre, white circles, Post, gray circles). *p < 0.05, ns: no significant difference

### Control of glutamate release by ChIs in D1- and D2-MSNs involves different mechanisms

Having shown that ChIs decrease the release of glutamate from presynaptic terminals synapsing on both D1- and D2-MSNs, we questioned whether these two neuronal populations shared the same mechanisms. There is strong evidence that ChIs induce DA release in the striatum (17, 24). Because DA regulates glutamate release (25), the hypothesis that ChIs’ effects on glutamate release involved DA was tested in ChAT.ChR2.eYFP x DrD1-TdTomato mice. First, we confirmed that optogenetic stimulation of ChIs indeed evoked DA release (Fig. 4Ai, ii) measured with fast-scan cycling voltammetry (FSCV) when using the same stimulation pattern as that of our electrophysiological experiments. We then tested the putative role of DA in mediating ChIs-induced inhibition of glutamate release in D1-MSNs. ChI stimulation resulted in significantly longer IEIs in of D1-MSNs (Fig. 4B ACSF group as already presented in Fig 2G). In the presence of dopamine D1- and D2-receptor antagonists IEIs were no longer lengthened by optogenetic ChI stimulation (Fig. 4B, ACSF vs Sulp+SCH, F(1,2514) = 32.12, p<0.0001, MM GLM), with Tukey post-hoc tests revealing significant differences between Pre and Post conditions in ACSF condition, while the Pre- and Post-stimulation groups with dopamine antagonists were not significantly different. These findings show that the effect of ChI activity is DA-dependent in D1- MSNs with DA released by ChI stimulation likely acting on DA receptors expressed on glutamatergic terminals (26, 27). Next, the dependency of the DA effect on nAChR activation was confirmed (Fig. 4B, ACSF vs Mec, F(1,2562) = 28.25, p<0.0001, MM GLM, with Tukey post-hoc tests showing no differences between Pre and Post conditions when nAChRs were blocked). Antagonizing mAChR signaling also prevented the ChI-activation mediated increase in IEI (Fig. 4B, ACSF vs Atr, F(1,2269) = 56.81, p<0.0001, MM GLM, with Tukey post-hoc tests showing no differences between Atr Pre and Post conditions). In line with the preceding outcomes, antagonists of both mAChR and nAChR also blocked the increase of IEIs (Fig. 4B, ACSF vs Atr+Mec, F(1,2521) = 28.15, p <0.0001, MM GLM, with Tukey post-hoc tests showing no differences between Atr+Mec Pre and Post conditions). Finally, bath application of antagonists did not significantly change glutamate release under baseline conditions (Fig. S3 D1-MSNs, F(4,60) = 1.472, p = 0.22, one-way ANOVA). Together, these results indicate that mAChR, nAChR and DA receptor signaling are all required to mediate the effects of ChIs on glutamate transmission in D1-MSNs.

**Figure 4.**
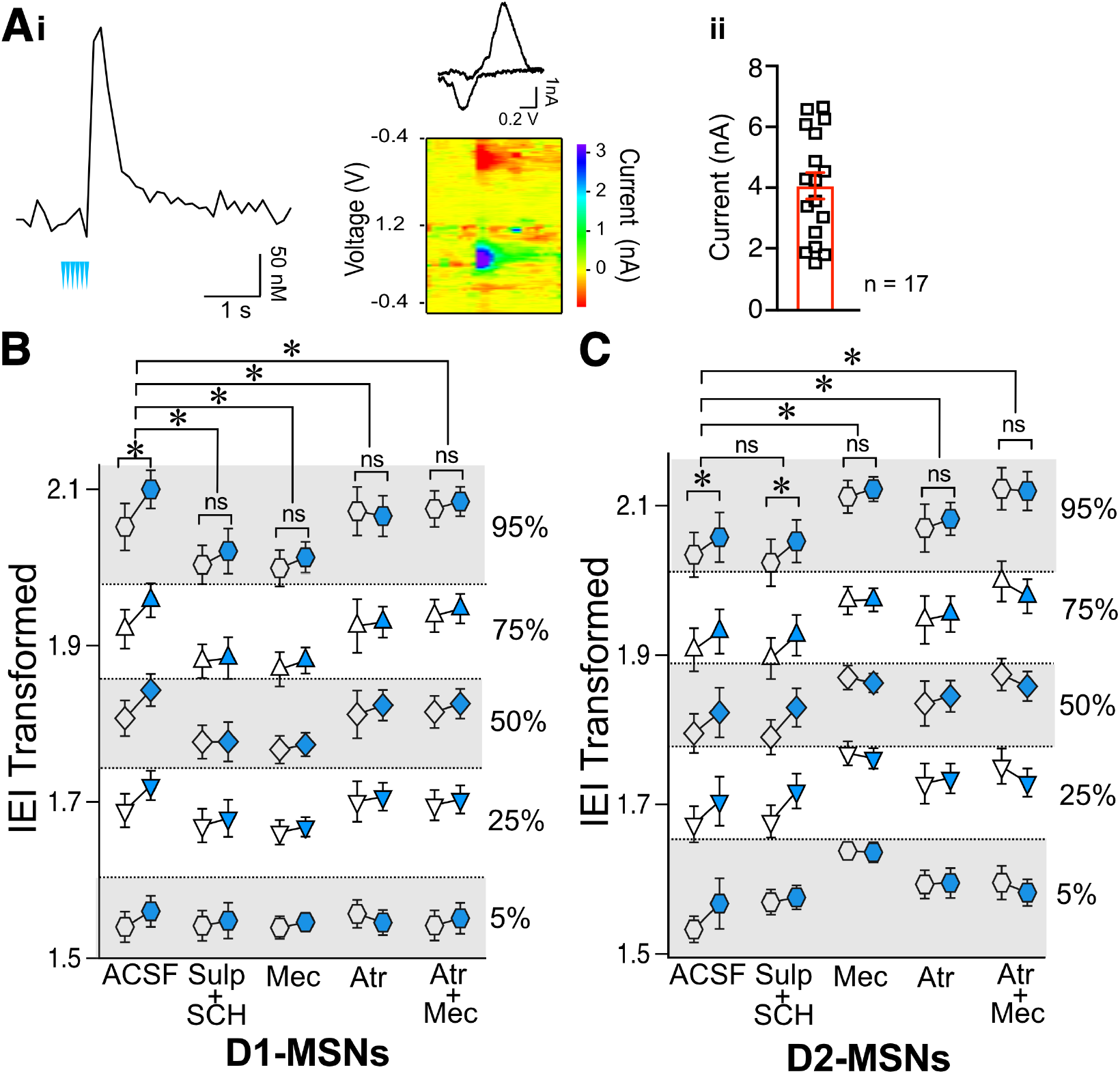
Effects of ChIs on D1- and D2-MSNs glutamate release are mediated through different pathways. **A**. Optogenetic activation of ChIs evokes dopamine release in NAc as measured by voltammetry. **i**. Representative DA trace and cyclic voltammogram showing characteristic DA waveform. **ii**. DA responses evoked from ChI stimulation scatter plot and average <u>±</u> SEM showing the range of DA currents from 6 slices and 17 recordings. **B**. Transformed IEI of D1-MSNs sEPSCs shown as percentiles of cumulative distribution. Data for the control group (ACSF) is reproduced from Fig. 2G. IEI is shown before (Pre, white circles) and after (Post, blue circles) ChIs stimulation in control conditions (ACSF) and in presence of antagonists: Sul+SCH (5 uM SCH-23390, D1 receptor antagonist + 1 uM sulpiride, D2 receptor antagonist), Atr (1 uM atropine, mAChR antagonist), Mec (5 uM mecamylamine, nAChR antagonist) and Atr+Mec (atropine + mecamylamine). **C:** Same as B in D2-MSNs. *p < 0.05, ns: no significant difference

Next, the mechanisms underlying ChIs-mediated inhibition of glutamate release were assessed in D2-MSNs. Surprisingly, in D2-MSNs the effect of ChI stimulation on IEI length was not significantly different between the ACSF and DA receptor antagonists groups (Fig. 4C, ACSF vs Sul-SCH, F(1,1821) = 0.2625, p = 0.6085, MM GLM), with ChI stimulation significantly lengthening IEI in both groups (p < 0.05). This suggested that ChIs exert a direct presynaptic control over glutamate release in D2-MSNs that does not require DA. Interestingly, blocking nAChR signaling in D2-MSNs prevented the ChI stimulation induced lengthening of IEI (Fig. 4C ACSF vs Mec, F(1,1804) = 25.34, p<0.0001, MM GLM, with Tukey post-hoc tests showing no difference in Mec Pre and Post groups). Recordings in the presence of the mAChR blocker atropine were also significantly different from ACSF control (Fig. 4C ACSF vs Atr, F(1,1753) = 3.537, p = 0.0402, MM GLM, with Tukey post-hoc tests revealing no difference in the Atr Pre and Post groups). Accordingly, simultaneous application of both nAChR and mAChR antagonists found similar blockage of the ChI effect on IEI (Fig. 4C, ACSF vs Atr+Mec, F(1, 1760) = 55.05, p<0.0001, MM GLM, with Tukey post-hoc tests showing no difference in Atr+Mec Pre vs Post groups). These findings indicate that, unlike in D1-MSNs, ChI-mediated decrease in glutamate release in D2-MSNs is DA-independent but still mediated by both mAChR and nAChR antagonists.

While muscarinic and DA receptors antagonists applied under baseline conditions (without optogenetic stimulation) did not alter sEPSCs IEIs (Fig. S3, D2-MSNs, Atr: p = 0.29, Atr+Mec: p = 0.11, Sul+SCH: p = 0.20), blocking nicotine receptors with mecamylamine significantly increased IEIs (Fig. S3, D2-MSNs Mec, Mann-Whiteny U = 21, p = 0.0387, Mann-Whitney test). This finding provides a possible explanation for the observation that blocking nicotinic receptor with mecamylamine in D2-MSNs increased IEI: bath application of mecamylamine significantly decreased glutamate release, thus reaching a “floor effect” that could not be further decreased by ChIs stimulation (Fig. 4C), indicating high nicotinic receptor sensitivity to the baseline tonic ACh release.

### Binge alcohol drinking selectively reverses the effect of ChI-mediated glutamatergic synaptic transmission in D1-MSNs

The effects of preceding alcohol exposure on the ChI control of MSN excitability was assessed using the drinking-in-the-dark (DID) paradigm, a well-established model of binge alcohol drinking (28). The DID paradigm allows mice to drink 20% alcohol for 2h starting 2h into the dark phase for 5 consecutive days per week (Fig. 5A). After 2 weeks of drinking either 20% alcohol (DID group) or water (Naïve group), sEPSCs were recorded in D1- and D2-MSNs before and after optogenetic stimulation. As previously shown in Figs. 2 and 3, we constructed cumulative distribution plots for D1-MSNs (Naïve, Fig. 5B and DID Fig. 5C) and D2-MSNs (Naïve, Fig. 5D and DID Fig. 5E). Suprisingly, unlike in Naïve conditions, the effect of alcohol exposure on ChI regulation of sEPSC IEI length was significantly different in D1- and D2-MSNs (Fig. 5F, F(1,4778) = 12.08, p = 0.0005, MM GLM, 3-way interaction between optogenetic treatment, alcohol treatment and cell type). Tukey post-hoc tests revealed that in naïve D1-MSNs, ChI activation increased IEI while in DID exposed D1-MSNs, IEIs were significantly decreased, thus reversing the ChI effect on glutamatergic transmission. Conversely, ChI activation resulted in longer IEIs in D2-MSNs of both naïve and DID exposed mice. Interestingly, the effects of alcohol treatment seen in D1-MSNs depended on the size of IEI (Fig. 5F D1-MSNs, F(4,2374) = 3.900, p = 0.0037), with post-hoc tests revealing differences in the 75 and 95 percentiles, but not other intervals, indicating that alcohol effect was especially pronounced in largest size IEIs. In D2-MSNs there was no relationship between IEI size the effect of alcohol. These results demonstrate that preceding alcohol exposure selectively inverts the effect of ChI activation on D1-MSNs from reducing to increasing glutamate release onto D1-MSNs, while not having this effect on D2-MSNs. There was no significant difference in DID D1-MSNs sEPSC amplitude (Fig. S4A, t(13) = 1.46, p = 0.1680, paired t-test) or in DID D2-MSNs sEPSC amplitude (Fig. S4B, t(10) = 0.6948, p = 0.503, paired t-test).

**Figure 5:**
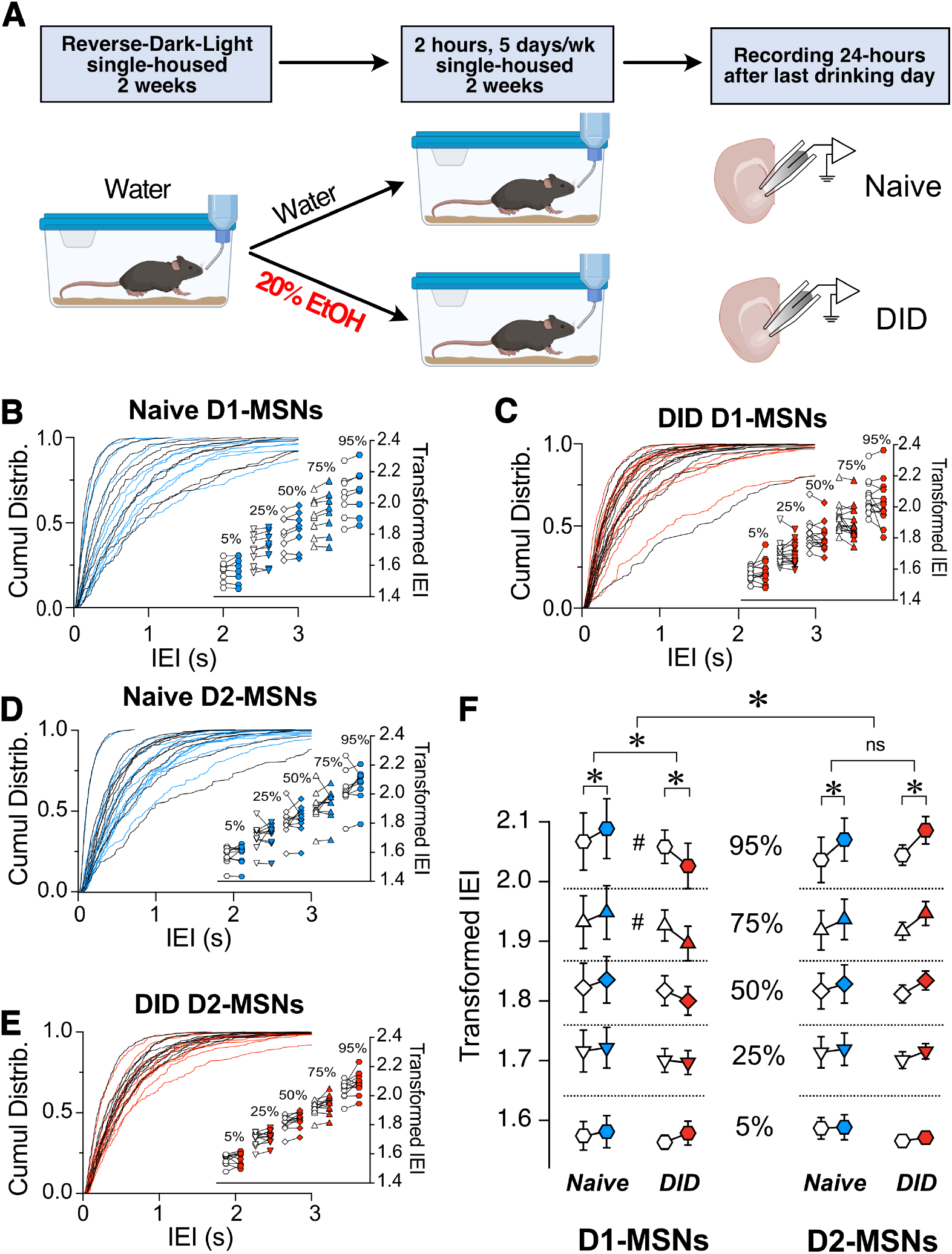
Binge alcohol drinking differentially affects the control by ChI on glutamate release in D1 and D2R MSNs. **A**. Schematic of drinking in the dark (DID) treatment. Mice were single-housed at 4-5 weeks of age and placed into reversed dark-light schedule to habituate for 2 weeks and then given either water (Naïve group) or 20% EtOH (DID group) every day for 2 hours, 5 days a week for 2 weeks. Brain slices were then isolated, and MSNs’ sEPSCs recorded. **B:** Cumulative frequency distribution of D1-MSN IEI of sEPSCs in ChAT-ChR2 mice group before (Pre, black traces) and after (Post, blue traces) ChI optogenetic stimulation. Each solid line represents a neuron. Average traces for Pre and Post conditions are shown in dotted black and blues lines, respectively. **Inset**. Cumulative distributions of D1-MSNs EPSCs IEIs in ChAT-ChR2 mice broken into percentiles of distribution to quantify median (50%), shoulders (25 and 75%), and extreme values (5 and 94%) of distribution, that are 1^10 transformed to normalize the distribution.**C**. Same as B in DID mice. Black and red traces indicate IEI before and after ChI, respectively. (n = 14). **D:** Cumulative frequency distribution of D2-MSN IEI of sEPSCs in ChAT- ChR2 mice before (Pre, black traces) and after (Post, blue traces) ChI optogenetic stimulation. Each solid line represents a neuron. Average traces are shown in dotted black and blues lines for Pre and Post conditions, respectively. **Inset**. Cumulative distributions of ChAT-ChR2 D2-MSNs EPSCs IEIs broken into percentiles of distribution to quantify median (50%), shoulders (25 and 75%), and extreme values (5 and 94%) of distribution, that are 1^10 transformed to normalize the distribution. (n = 10). **E:** Same as B in DID mice. Black and red traces indicate IEI before and after ChI, respectively. (n = 11). **F**. Percentiles of cumulative distribution of transformed IEIs of EPSCs in Naïve ChAT-ChR2 (Pre, white circles, Post, blue circles) and DID ChAT-ChR2 (Pre, white circles, Post, red circles) in D1- and D2-MSNs. *p < 0.05, #p < 0.05, ns: no significant difference

Given the effects of preceding alcohol exposure on ChI regulation of glutamate release in D1-MSNs, the next objective was to identify receptors mediating these effects. Interestingly, in DID-exposed D1-MSNs, recording in the presence of D1- and D2-antagonists seemed to block the effect of DID (Fig. 6 ACSF vs Sulp+SCH, F(1,2331) = 40.49, p < 0.0001, MM GLM). This increase of IEI length following ChI stimulation in the presence of dopamine receptor antagonists in DID-exposed D1-MSNs was reminiscent of the naïve D1-MSN group (Fig. 5F), indicating an important role of DA receptors in alcohol’s effect on ChI-modulated glutamate release. In the presence of mAChR antagonist atropine, ChI effect of sEPSC IEI was also significantly different from ACSF condition (Fig. 6 ACSF vs Atr, F(1,2495) = 25.1129, p < 0.0001, MM GLM, with Tukey post hoc showing a significant IEI decrease only in ACSF group, but no significant difference between Pre vs Post groups in the Atr group). Finally, recording in the presence of nAChR antagonist mecamylamine also abolished the effect of ChIs, and was significantly different from ACSF (Fig. 6 ACSF vs Mec, F(1,2577) = 6.281, p = 0.0123, MM GLM, with Tukey Post hoc similarly showing only a significant difference in the ACSF group). The most parsimonious interpretation of these results is that the influence of preceding alcohol exposure on the ChI-mediated glutamate release in D1-MSNs mostly depends on dopaminergic signaling with additional influences from both nAChR and mAChR signals.

**Figure 6.**
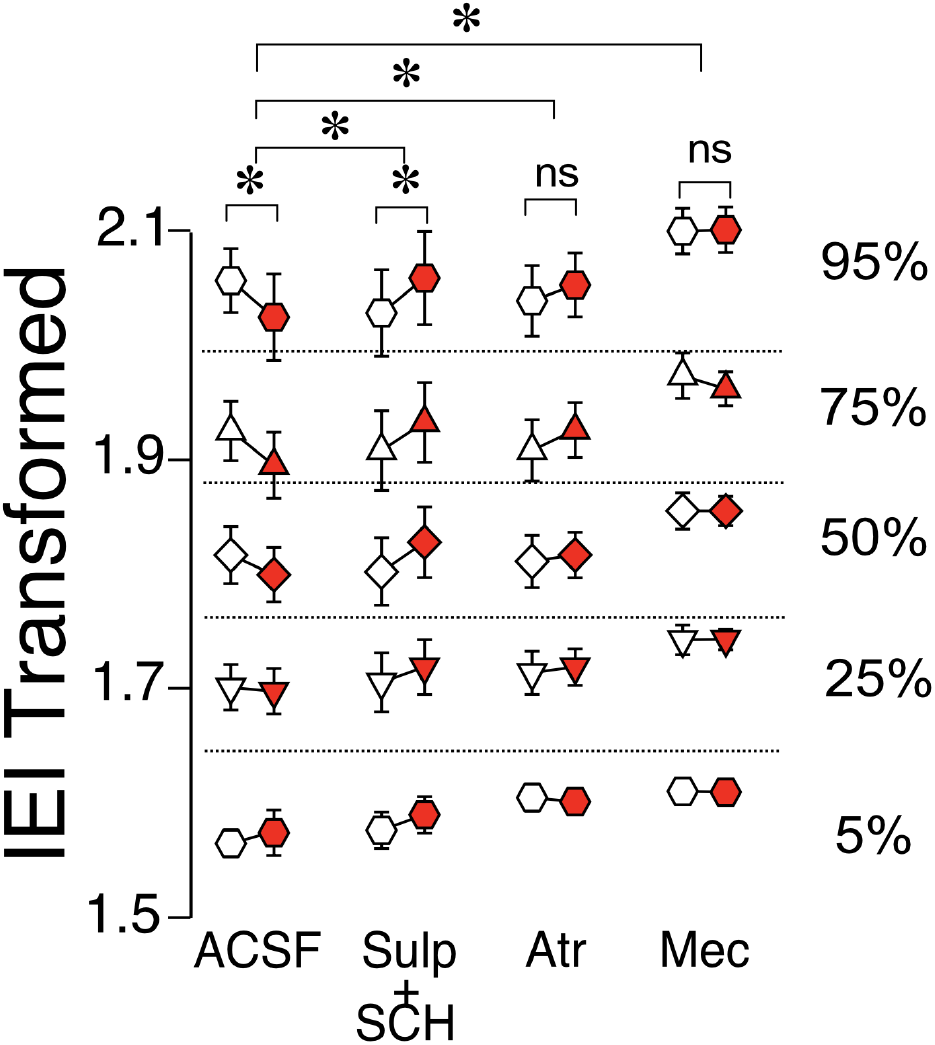
Effects of ChI stimulation in DID D1-MSNs in the presence of antagonists. IEI in DID D1-MSNs EPSCs shown as percentiles of cumulative distribution. Data for the control group (ACSF) is reproduced from Fig. 5F. Each group of Pre (white circles) and Post (red circles) column is recorded either without (ACSF) or with the bath presence of antagonists: Sul+SCH (5 uM SCH- 23390, D1 receptor antagonist + 1 uM sulpiride, D2 receptor antagonist), Atr (1 uM atropine, mAChR antagonist), Mec (5 uM mecamylamine, nAChR antagonist). *p<0.05, ns: no significant difference

Interestingly, the application of bath antagonists in DID D1-MSNs changed only during mecamylamine treatment compared to ACSF (Fig. S5, Mec, Mann-Whiteny U = 22, p = 0.0065), while the other antagonists were not different from ACSF group (Fig. S5, Atr: p = 0.693, Sul+SCH: p = 0.896). This is in stark contrast from naïve D1-MSN group (Fig. 4B), where mecamylamine treatment did not change the baseline, but is reminiscent of naïve D2-MSN group (Fig. 4C), where mecamylamine also increased IEIs.

### ChI optogenetic stimulation in vivo in the NAc increases alcohol consumption in mice

Since binge alcohol drinking modulates the ChI-mediated synaptic transmission onto MSNs *ex vivo*, we next tested whether optogenetic stimulation of ChIs in freely-moving animals altered alcohol consumption. Fiber optic cannulas were implanted into the NAc of 4-5 weeks old mice that were allowed to recover and habituate to our reverse-light-dark room for 3 weeks before being optogenetically stimulated during the first 4 days of alcohol exposure (Figs. 7A and B). From the very first day, the volume of alcohol consumed by stimulated ChAT.ChR2 mice group (Fig. 7C, ChR2) was markedly larger compared to mice in the non-stimulated ChR2 group (Fig. 7C, ChR2 NS) and stimulated ChAT.Cre controls (Fig. 7C, F(2, 22) = 7.685, p = 0.0029, MM GLM, with Tukey post-hoc tests showing ChR2 group significantly different from both control groups). The pattern of alcohol consumption was determined using lickometers by measuring the number and timing of licks of the drinking spout delivering alcohol (Fig. 7D). The total number of licks in stimulated ChR2 mice during the 4 days was significantly increased compared to the non-stimulated and Cre control groups (Fig. 7E, F(2,22) = 6.50, p = 0.0061, MM GLM). The increase in alcohol drinking was likely due to the increased frequency of consumption measured as the licking bout IEI was dramatically reduced in the ChR2 group (Fig. 7F, F(2,20) = 5.376, p = 0.0135, one-way ANOVA), and licks were highly correlated with the amount of alcohol consumed (Fig S6A). General locomotor activity of a subgroup of mice was measured using a passive infrared activity monitoring system and compared to alcohol licks during the same time interval (Figs. 7G, H). This comparison illustrated that the increased drinking in the stimulated ChR2 mice could not be explained through a general increase in activity levels in these mice (Fig. 7H; ChR2: R^2^ = 0.134, p = 0.122; Cre controls: R^2^ = 0.023, p = 0.472). The increased alcohol consumption observed in stimulated ChR2 mice was specific to alcohol consumption and did not extend to the consumption of saccharine (Fig. 7I, F(1,47) = 1.53, p = 0.22, RM two-way ANOVA, Fig. S6B, F(1,47) = 1.78, p = 0.93, RM two-way ANOVA), and water (Fig. 7J, F(1,45) = 0.33, p = 0.57, RM two-way ANOVA, Fig. S6C, F(1,49) = 0.03, p = 0.87, RM two-way ANOVA). Finally, stimulated ChR2 and Cre control mice (Fig. 7K) did not differ in ambulation time course (F(7,84) = 0.80, p = 0.59, RM two-way ANOVA), total ambulation (t(12) = 1.63, p = 0.13, student’s t-test), fine movement (t(12) = 1.12, p = 0.28, student’s t-test), or vertical motion (rearing, Fig. S6D, t(12) = 0.8, p = 0.44, student’s t-test). These results demonstrate that optogenetic stimulation of ChIs specifically altered alcohol consumption, without affecting water and saccharine drinking, an effect that was not due to increased activity levels.

**Figure 7:**
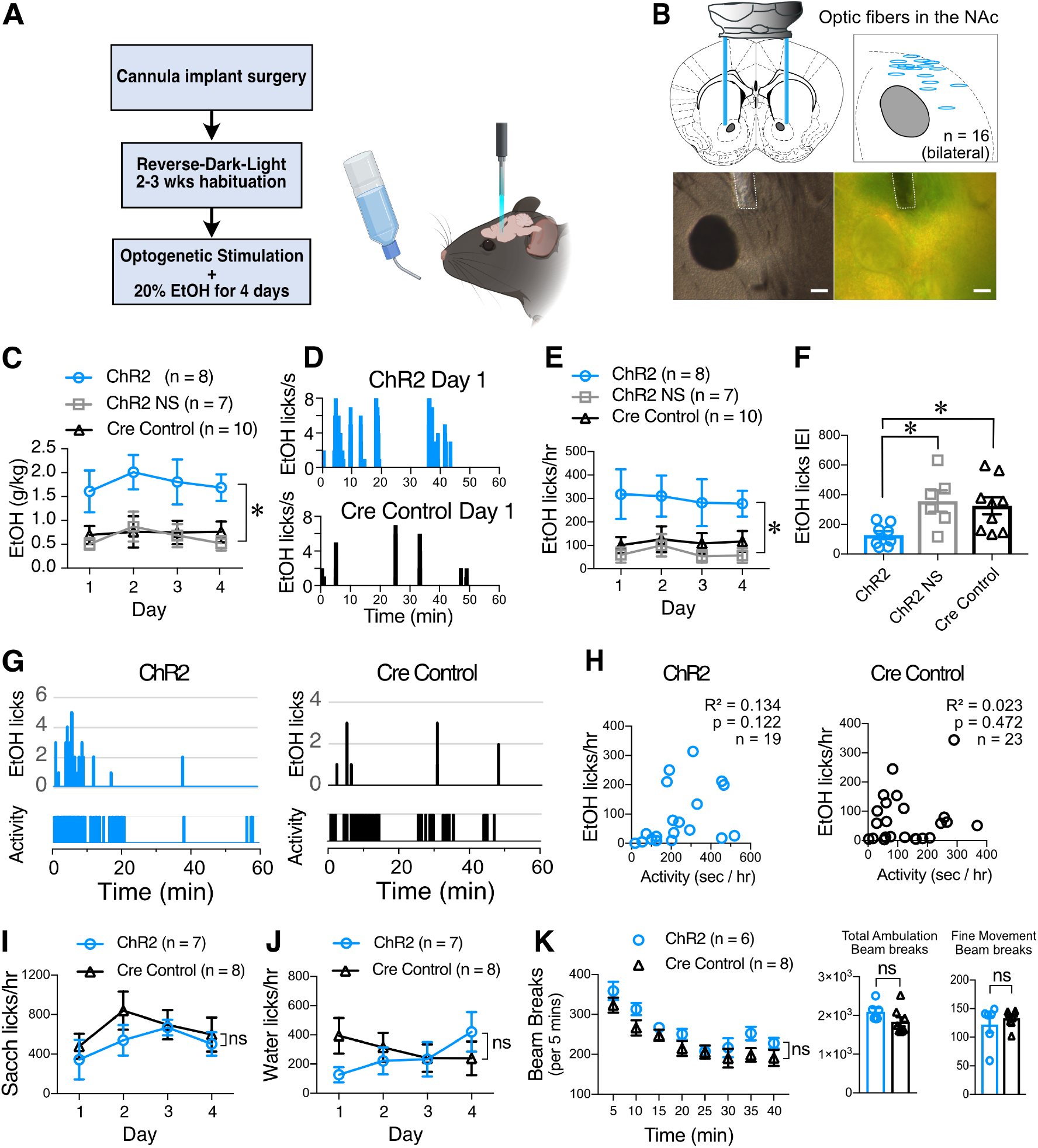
ChI optogenetic stimulation increases EtOH consumption. **A**. Schematic of the DID behavioral experiment. ChAT.ChR2 and ChAT.cre mice underwent fiber optic implant in the NAc surgery at 4-5 weeks of age, recovered and habituated to reverse-dark-light schedule. ChAT.ChR2 and ChAT.cre mice were optogenetically stimulated during 4 days of 20% alcohol exposure and tethered to the fiber cord while ChAT.ChR2 not stimulated (NS) mice were only tethered to the fiber cord. **B**. Schematic of the bilateral optic fiber cannula implant in the NAc (top left panel), locations of bilateral implants in 8 ChR2 mice that were used for the experiment (top right panel), example image of cannula placement in DIC (bottom left panel) and fluorescent light (bottom right panel). **C**. Average daily alcohol consumption over 4 days in optically stimulated ChR2 mice (ChR2), non-stimulated ChR2 mice (ChR2 NS) and stimulated ChAT-cre mice (Cre controls). **D**. Graphs of alcohol licks during 1st hour period in representative stimulated (top graph, blue bars) and non-stimulated ChR2 mice (bottom, gray bars). **E**. Daily average of alcohol licks over 4 days of alcohol exposure in optically stimulated ChR2 mice (ChR2), non-stimulated ChR2 mice (ChR2 NS) and stimulated ChAT-cre mice (Cre controls). **F**. Frequency of alcohol licks as measured by average licking bout inter-event interval (IEI) per mouse over a 4-day period. **G**. Representative plots of mouse activity as measured by passive infrared (PIR) activity monitoring system and corresponding EtOH licks in ChR2 (Left graphs, blue bars) and Cre control mice (right graphs, black bars). **H**. The number of licks is not correlated with mice locomotor activity in ChR2 (blue symbols) and Cre control groups (black symbols). **I**. Saccharine consumption measured as number of licks/hr over 4 days in 1 hr-long optogenetically stimulated ChR2 (blue circles) and Cre control mice (black triangles). **J**. Water consumption measured as number of licks/hr over 4 days in 1 hr- long optogenetically stimulated ChR2 and Cre control mice. **K**. Ambulatory activity test of ChR2 and Cre control mice during ChI optogenetic stimulation. Ambulation time course shows average beam breaks every 5 mins for 40 mins of the test. Total Ambulation shows the total beam breaks in 40 min, fine movement shows grooming activity and vertical ambulation shows rearing activity. *p < 0.05, ns: no significant difference.

## Discussion

The output neurons of the NAc, D1- and D2-MSNs, are a key part of the neurobiological mechanisms underlying drug addiction (7) and altering their outputs will likely be an important part of any future treatments of alcohol addiction (29). ChIs provide a promising avenue to do so since although these cells make up only 1-2% of the NAc neuronal population, they fulfill a key integrative role modulating the activity of MSNs(5). The data presented here show that ChIs control glutamatergic synaptic transmission in both D1- and D2-MSNs in alcohol-naïve mice, though the underlying regulatory mechanisms differ (Naïve, Fig. 8). While the ChIs-driven decrease of glutamate release onto D1-MSNs is mediated by nicotinic and muscarinic ACh receptors through DA receptors, ChIs control of glutamate release onto D2-MSNs likely stems from ChIs directly synapsing on glutamatergic terminals through nicotinic and muscarinic ACh signals. Surprisingly, preceding alcohol exposure results in a switch where the effect of ChIs activity inverts from inhibiting to potentiating glutamatergic transmission in D1-MSNs while their inhibitory effect in D2-MSNs remains unchanged (Fig. 8). Based on this dramatic change of its influence on D1-MSNs we hypothesized that altering ChI activity could be used to modulate alcohol drinking behavior. In line with this hypothesis, ChI optogenetic stimulation *in vivo* increased alcohol consumption in mice without altering locomotor activity, saccharine or water consumption. Together, these findings identify NAc ChIs as key modulators of D1- and D2-MSNs excitability and suggest this cell population as a promising target of future addiction treatment strategies.

**Figure 8:**
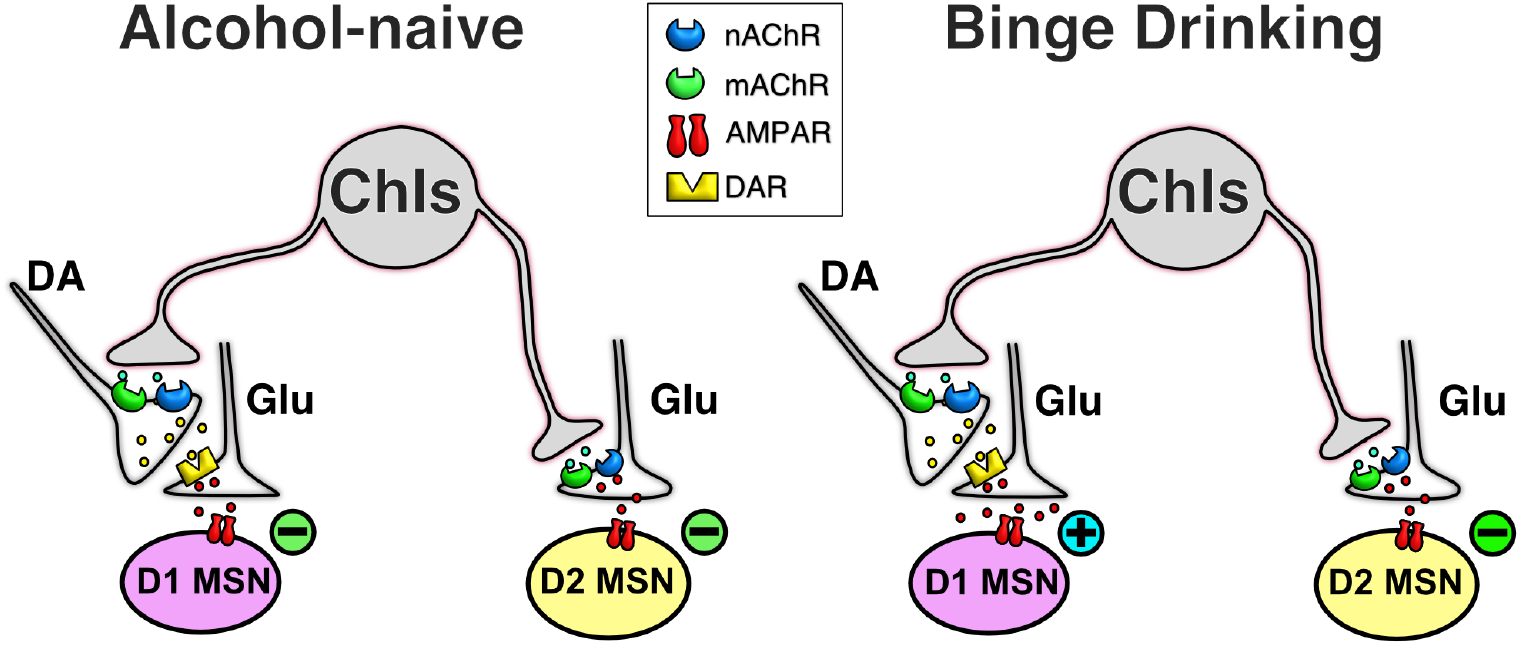
Simplified schematic of ChI effect on glutamatergic neurotransmission in MSNs in naïve and alcohol binge drinking mice. In D1-MSNs of naïve mice, ChIs decrease glutamatergic neurotransmission presynaptically through the release of DA in D1-MSNs (top left cartoon), an effect that is reversed in DID mice (top right cartoon). In contrast, in D2-MSNs of naïve mice, ChIs directly control glutamate release from terminals synapsing on D2-MSNs (bottom left cartoon), an effect that is unchanged in DID mice (bottom right cartoon)

### ChIs decrease glutamate release in D1- and D2-MSNs through different mechanisms

Our finding that ChIs inhibit MSN glutamatergic synaptic transmission through a presynaptic mechanism confirms previous reports showing that acetylcholine receptor (AChR) agonists reduce the probability of glutamate release in the striatum (30–33). Similarly, direct stimulation of ChIs depressed electrically evoked EPSCs, an effect also attributed to presynaptic cholinergic receptors (34, 35), although a postsynaptic effect was also reported (36). Despite decades of research striving to understand how ChIs regulate glutamatergic synaptic transmission in MSNs, the mechanisms mediating their effects on D1- and D2-MSNs glutamatergic synaptic transmission is still poorly understood (37, 38). Our study provides evidence that ChIs employ different mechanisms to regulate glutamate release in D1- and D2-MSNs. Specifically, our data demonstrates the influence of DA on ChIs’ regulation of glutamate release in D1- but not D2-MSNs. The role of DA is supported by data from several groups showing that ChIs evokes DA release (17, 24, 39, 40), likely through α*β2 nAChR expressed on DA terminals (17, 41). In addition to nAChRs, we found that mAChRs also contribute to the ChIs-mediated decrease of glutamate release in D1-MSNs, possibly through M5 mAChR (39, 42–44), a finding confirming previous studies (45–47). Our study also indicates that, upon its release, DA binds to DA receptors located presynaptically on glutamatergic terminals where they decrease glutamate neurotransmission, a finding supported by several studies (26, 48–50), likely by promoting adenosine efflux via A1 adenosine receptors (A1Rs) (51, 52). While our study confirms the role of DA and ACh receptors in regulating glutamatergic synaptic transmission, it provides key additional information as to how these neurotransmitters work together to regulate glutamate release in D1-MSNs.

As opposed to D1-MSNs, our data supports the notion that DA receptors do not contribute to ChI-mediated decrease of glutamate release in D2-MSNs. Instead, ChIs appear to send direct projections to glutamatergic terminals synapsing on D2-MSNs, an effect that our pharmacological experiments suggest is mediated by mAChRs. Although performed in conditions that did not differentiate D1- from D2-MSNs, several groups reported a similar contribution of mAChRs on glutamate release in the striatum (31, 36, 46, 47). Specifically, M2-4 mAChR located presynaptically on glutamatergic terminals directly decrease glutamate release by inhibition of P/Q-type VGCC and reduction of action potential–induced Ca^2+^ increases in the bouton (31, 36, 47). We have also found that application of 1 µM nicotinic antagonist mecamylamine decreases sEPSCs in D2-MSNs. However, much ambiguity still exists on the nicotinic effect on glutamate release. α4β2 nAChR antagonist has been shown to increase glutamate release (53), while 1 µM and 10 µM nicotine application was shown to decrease sEPSCs frequency (32), and 2 µM nicotine was also found to not change sEPSCs frequency (45). Finally, we have found that ChIs inhibited, albeit moderately (i.e., <10%), electrically evoked EPSPs in D2-MSNs only, mirroring a similarly small reduction of EPSCs in unidentified MSNs (35). These results emphasize the importance of distinguishing between striatal D1- and D2-MSNs when assessing their function in basal ganglia function.

### Alcohol exposure changes ChI control of the D1/D2 MSNs output balance

Unlike other drugs of abuse, alcohol does not have a single receptor, making identifying its targets difficult. Acute alcohol exposure modulates striatal output through ChIs (54) and inhibits ChIs firing (55), while chronic alcohol use reduces density of cholinergic varicosities (56). We found that ChIs’ stimulation increases alcohol consumption *in vivo*, while 2-week alcohol administration reverses the ChI control of glutamate release in D1-MSNs from inhibition to potentiation. Our finding is in line with previous studies showing that repeated exposure to alcohol potentiated D1- MSNs glutamatergic transmission (22, 29, 57, 58). In addition to increasing glutamate release from terminals synapsing on D1-MSNs, chronic alcohol exposure was shown to act postsynaptically by increasing of spines density in dendrites of NAc and dorsal striatum MSNs (59–61). Interestingly, glutamatergic transmission in D2-MSNs was not affected in binge alcohol drinking mice. Although this finding is somewhat surprising, Chen et al. reported that chronic alcohol exposure did not alter evoked EPSCs amplitude but increased GABAergic neurotransmission in dorsal striatal D2-MSNs (29). Although we can only speculate about the specific origin of NAc D2-MSNs inhibitory inputs, it is worth noting that MSNs are mostly inhibited by GABAergic interneurons that are under the control of ChIs (45, 62). If true, this would strengthen the putative central role that ChIs play in regulating excitability of D1- and D2-MSNs through glutamatergic and GABAergic synaptic transmission, respectively, and in shaping the overall message sent to downstream brain regions.

The mechanism responsible for reversing ChIs-mediated inhibition of glutamate release in D1- MSNs is unclear. Because DA is responsible for the ChIs-mediated decrease of glutamate release, increase of frequency observed in DID mice may result from alcohol either decreasing DA release (63) and/or impairing nAChR (64) and mAChRs function (65). Taken together, our findings offer a putative mechanism explaining why nAChR antagonists decrease alcohol consumption when administered i.p. (66–68), as well as directly into the NAc (69).

In summary, our study delineates a new understanding of the NAc circuitry and its effect on alcohol drinking behavior. ChIs likely induce DA release, which drives further alcohol consumption. Since ChI activation is what mediates this DA release, ChI stimulation *in vivo* will result in more DA released, driving the continuation of drinking after the very first sip (70). On the other hand, after 2 weeks of daily alcohol exposure, ChIs preferentially and repeatedly stimulate D1-MSNs, which leads to disbalance between D1- and D2-MSNs, potentiating the D1- MSNs “go” direct pathway and inhibiting the M2-MSNs “no-go” indirect pathway. ChI-mediated reciprocal strengthening of “go” and inhibition of “no-go” pathways could be a core element of compulsive increase of drinking over time and transitioning to addiction (1, 5). Therefore, inhibition of ChIs could be a future therapeutic target to treatment of alcohol use disorder.

## Supporting information

SFig. 1

SFig. 2

SFig. 3

SFig. 4

SFig. 5

SFig 6

## Disclosure

The authors have nothing to disclose.

**Supplemental Figure 1: D1-MSNs sEPSC and evoked EPSP measurements after ChI optogenetic stimulation. A**. Average sEPSCs amplitudes **before (**Pre, white circles) and after (Post, blue circles) ChI optogenetic stimulation in D1-MSNs ChAT.ChR2 mice (n = 14). **B**. Average sEPSCs amplitudes **before (**Pre, white circles) and after (Post, gray circles) light stimulation of ChIs in D1-MSNs TdTomato mice. (n = 10). **C**. Electrically-evoked EPSPs in ChAT-ChR2 D1-MSNs before (Pre, white circles), during (white circles, blue bar) and after (Post, blue circles) ChI optogenetic stimulation (n = 10). **D**. Electrically-evoked EPSPs in TdTomato control D1-MSNs before (Pre, white circles), during (white circles, blue bar) and after (Post, gray circles) ChI optogenetic stimulation (n = 7). *p<0.05, ns: no significant difference

**Supplemental Figure 2: D2-MSNs sEPSC and evoked EPSP measurements after ChI optogenetic stimulation. A**. Average sEPSCs amplitudes **before (**Pre, white circles) and after (Post, blue circles) ChI optogenetic stimulation in D2-MSNs of ChAT.ChR2 mice (n = 11). **B**. Average sEPSCs amplitudes **before (**Pre, white circles) and after (Post, gray circles) light stimulation in D2-MSNs of TdTomato mice. (n = 9). **C**. Electrically-evoked EPSPs in ChAT- ChR2 D2-MSNs before (Pre, black circles), during (Opto, white circles, blue bar) and after (Post, blue circles) ChI optogenetic stimulation (n = 9). **D**. Same as C in tdTomato mice (n = 7). *p<0.05, ns: no significant difference

**Supplemental Figure 3: sEPSCs IEI in the absence and presence of ACh and DA antagonists in D1- and D2-MSNs in naive mice**. Recordings of EPSCs IEIs during the bath application of antagonists. ACSF (control solution), D1 receptor antagonist Sulpiride + D2 receptor antagonist SCH-23390, nAChR antagonist mecamylamine, mAChR antagonist atropine. *p<0.05, ns: no significant difference

**Supplemental Figure 4: Binge alcohol drinking does not affect sEPSPs amplitude in D1- and D2-MSNs. A**. Amplitudes of DID D1-MSNs sEPSCs before (Pre, white circles) and after (Post, red circles) ChI optogenetic stimulation in ChAT.ChR2 mice (n = 14). Each symbol represents a MSN. **B:** Amplitudes of DID D2-MSNs sEPSCs before (Pre, white circless) and after (Post, red circles) ChI stimulation in ChAT.ChR2 mice (n = 11). Each symbol represents a MSN. *p<0.05, ns: no significant difference

**Supplemental Figure 5: sEPSCs IEIs in D1-MSNs in presence of ACh and DA receptors antagonists in DID mice**. Average sEPSCs IEIs in DID D1-MSNs recorded in ACSF (control solution) and in presence of dopamine D1 and D2 receptor (Sul+SCH), mAChR (Atrop), and nAChR (Mec) antagonists groups. *p<0.05, ns: no significant difference

**Supplemental Figure 6: ChI optogenetic stimulation and EtOH consumption behavior. A**. Graph shows strong correlation between the volume of EtOH consumed and the number of licks in ChAT.ChR2 (ChR2) and ChAT.cre mice (Cre control). **B**. 0.3% Saccharine consumed (g/kg) during 1-hr long ChI optogenetic stimulation in ChAT.ChR2 and ChAT.cre mice. **C**. Water consumed (g/kg) during 1 hr-long ChI optogenetic stimulation in ChAT.ChR2 and ChAT.cre mice. **D:** Vertical motion (rearing) in ChAT.ChR2 and ChAT.cre mice during ChI optogenetic stimulation. *p<0.05, ns: no significant difference

