## Supplementary figures and images for "Binge alcohol drinking alters the differential control of cholinergic interneurons over nucleus accumbens D1 and D2 medium spiny neurons"

### SFig 6

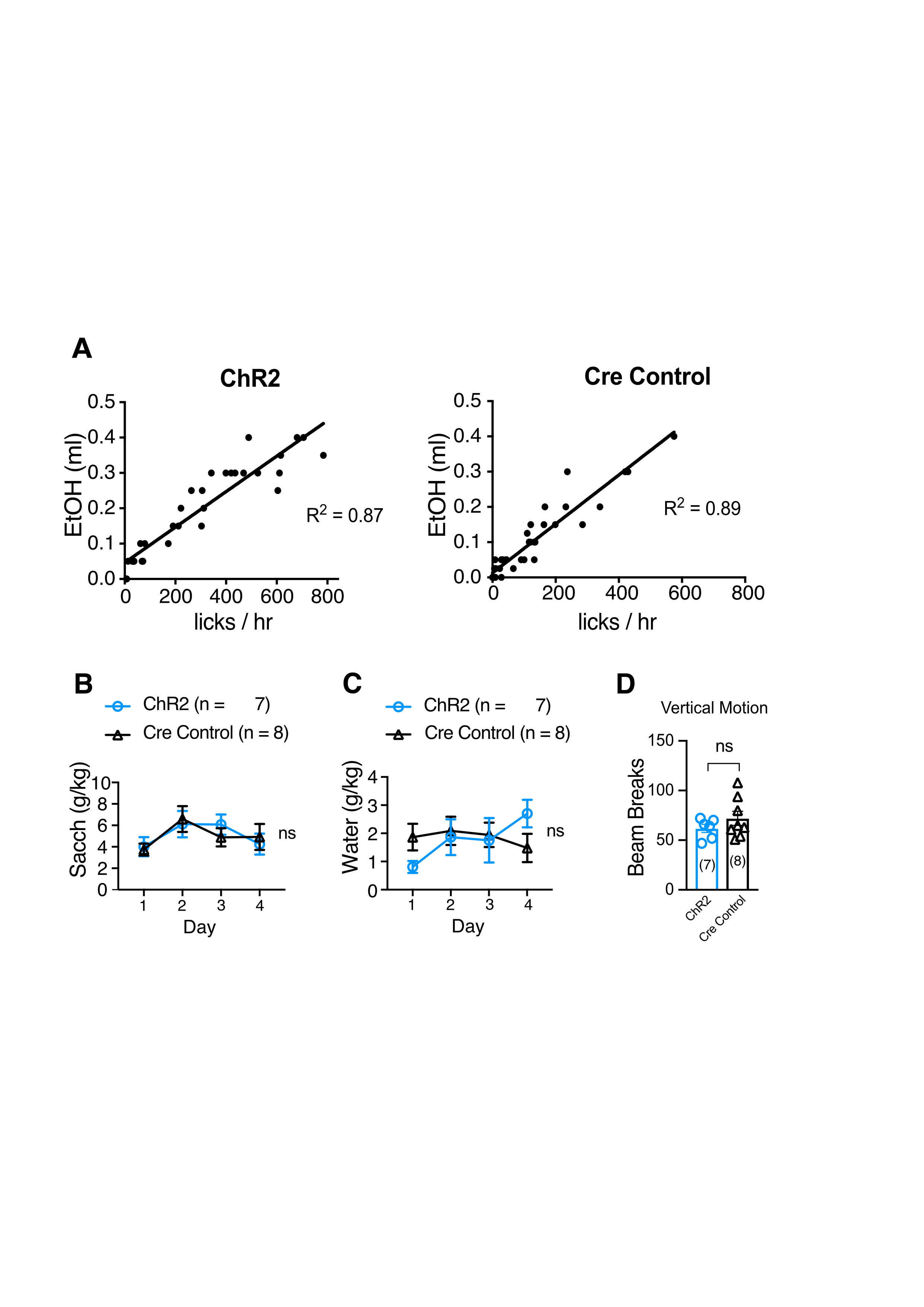

### SFig. 1

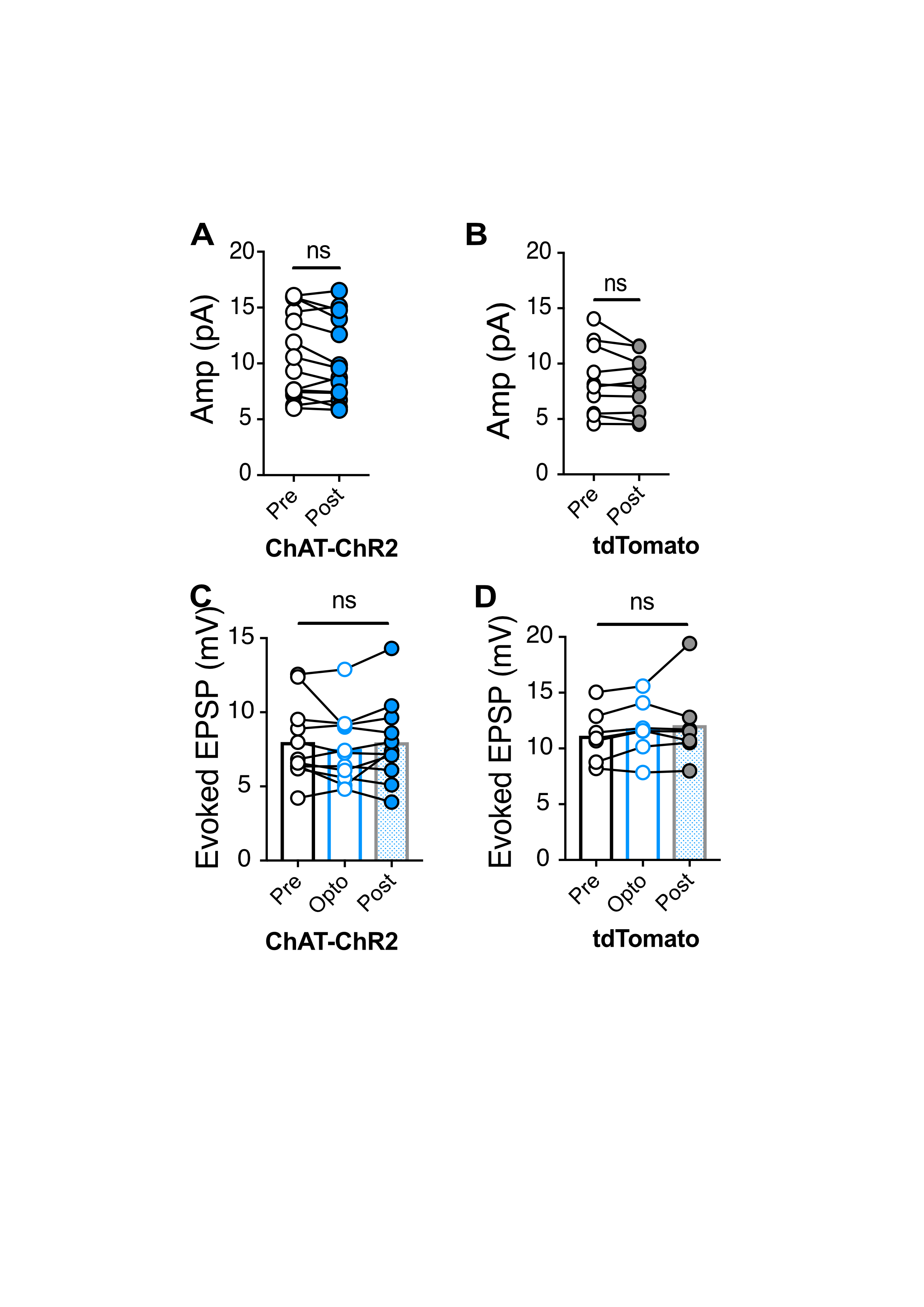

### SFig. 2

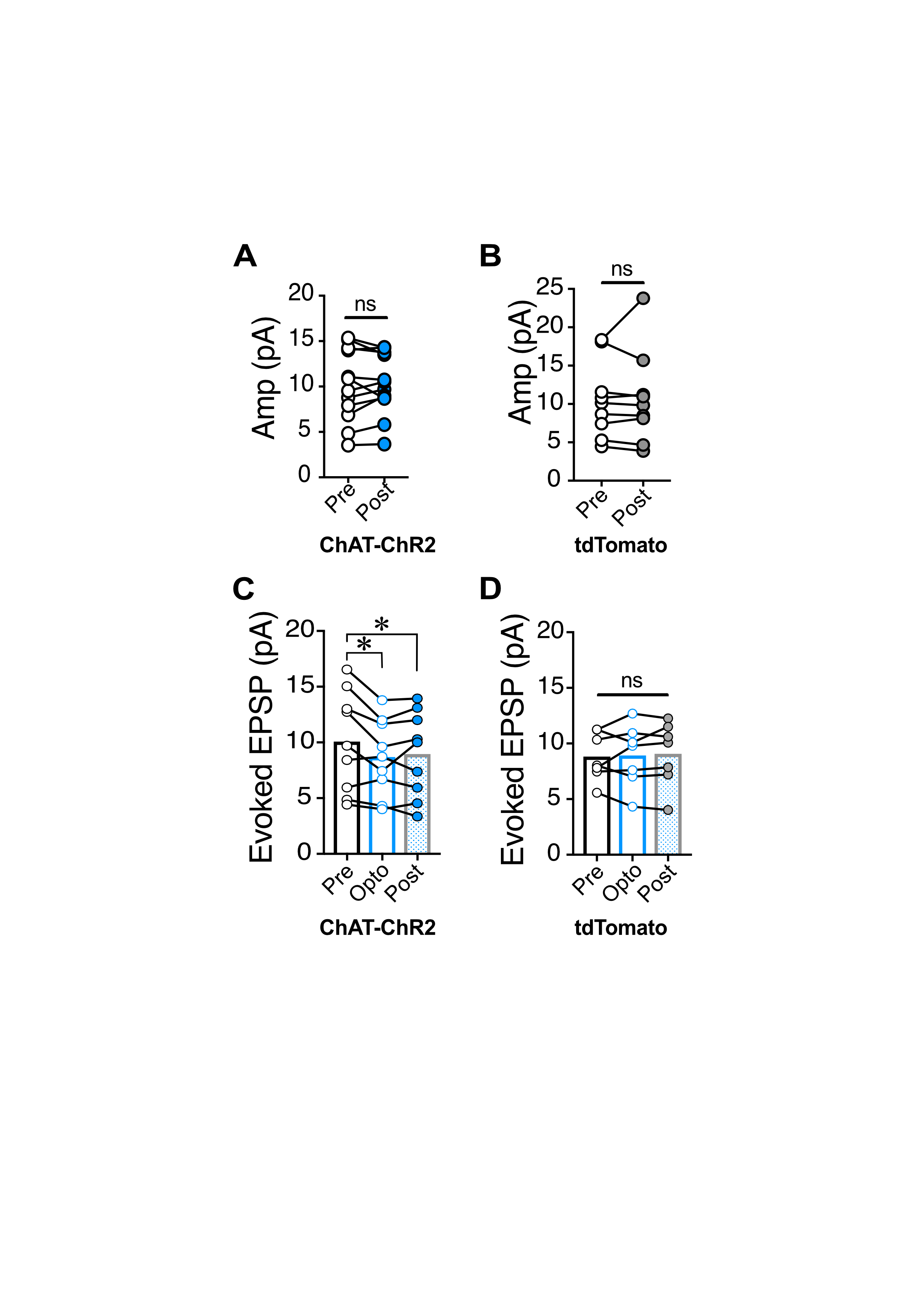

### SFig. 3

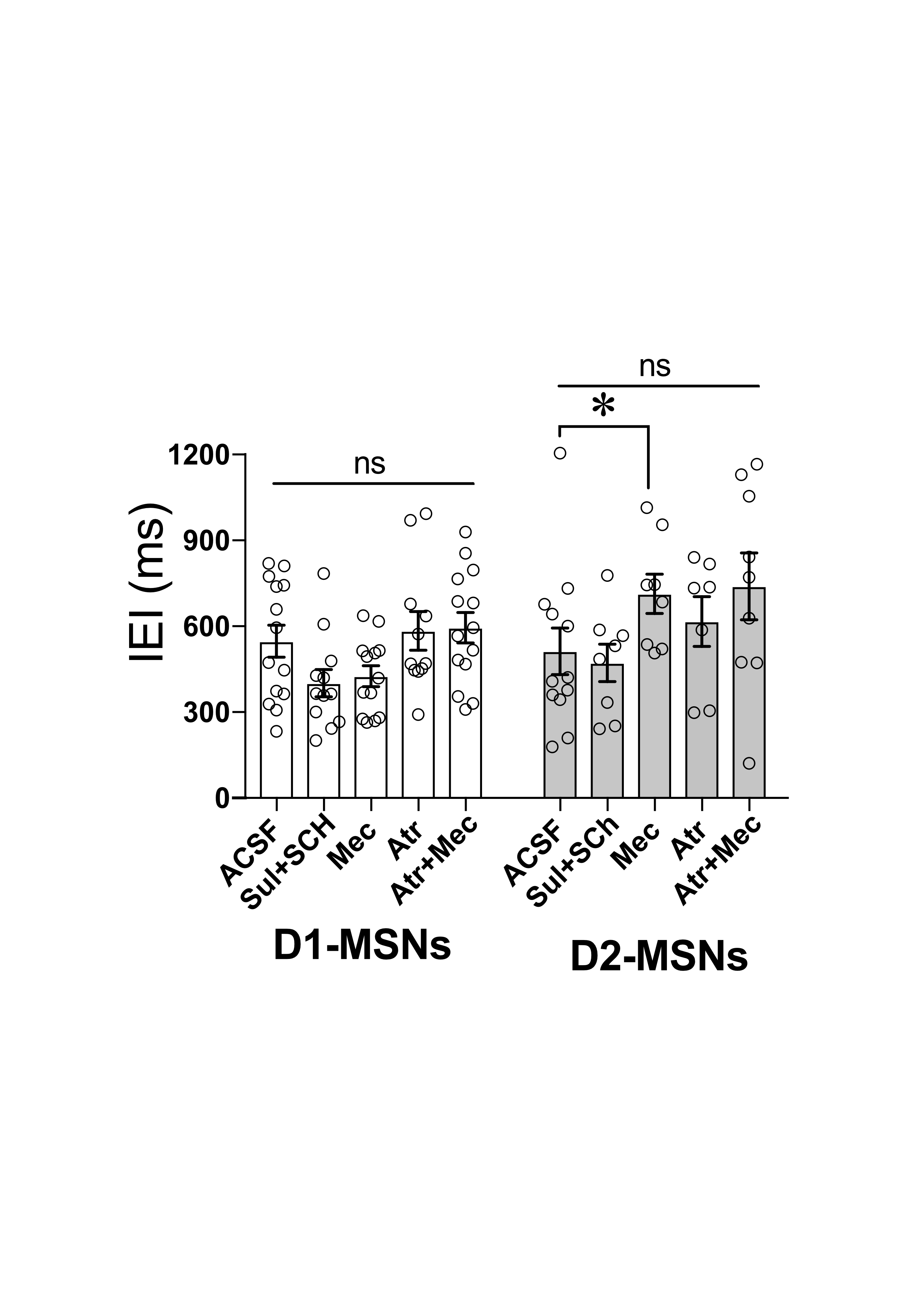

### SFig. 4

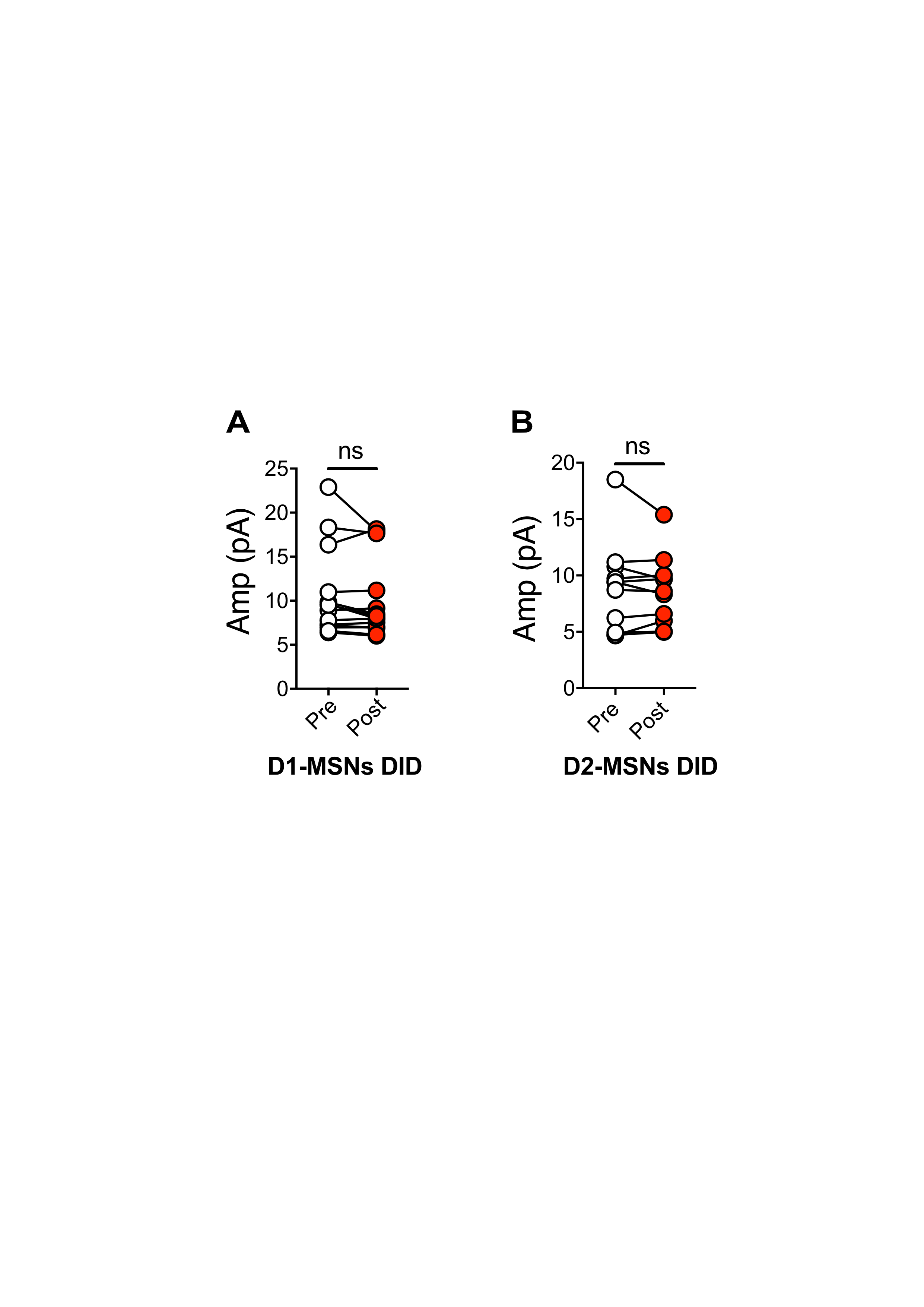

### SFig. 5

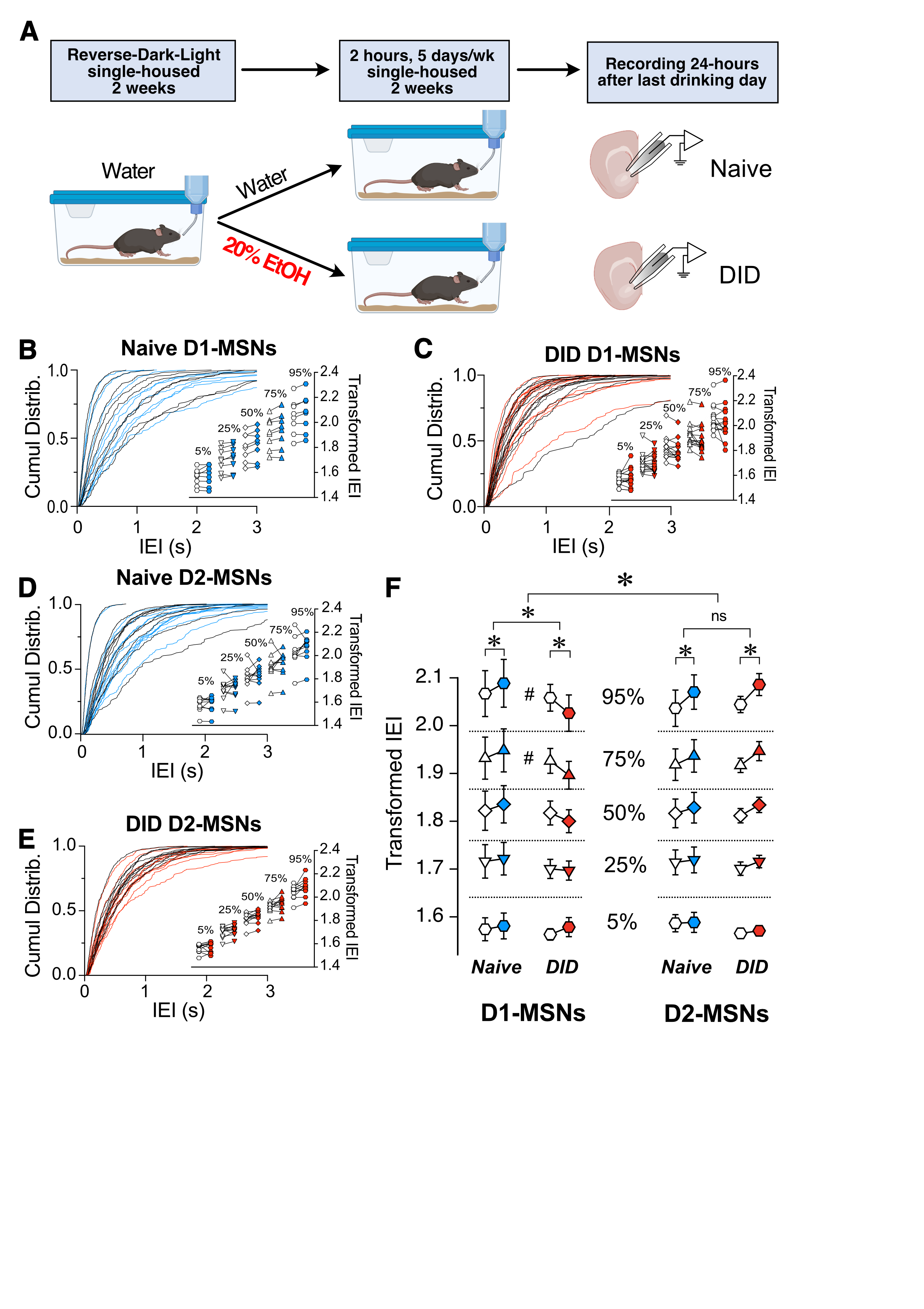
